# Analysis of cell death induction by the barley NLR immune receptor PBR1

**DOI:** 10.1101/2023.01.15.524147

**Authors:** Namrata Jaiswal, Ariana Myers, Terri L. Weese, Morgan E. Carter, Steven R. Scofield, Matthew Helm

## Abstract

The barley (*Hordeum vulgare* subsp. *vulgare*) disease resistance protein *Avr<u>P</u>ph<u>B R</u>esponse 1* (PBR1) mediates recognition of the *Pseudomonas syringae* effector, AvrPphB. PBR1 belongs to the coiled-coil nucleotide-binding leucine-rich repeat (CNL) family. However, little is known about the molecular mechanisms that lead to PBR1-dependent cell death (hypersensitive reaction; HR) in response to AvrPphB. Here, we investigated PBR1 immune signaling after *Agrobacterium*-mediated transient expression in *Nicotiana benthamiana*. We found that co-expression of PBR1 with AvrPphB resulted in robust cell death, confirming previous observations that PBR1 is indeed the cognate NLR that recognizes AvrPphB. The N-terminal tagging of PBR1 with super Yellow Fluorescent Protein (sYFP) abolished PBR1-mediated cell death, demonstrating that an N-terminal epitope tag disrupts PBR1-mediated immune signaling. Furthermore, none of the individual protein domains or truncations of PBR1 induced a HR-like cell death response as strong as full-length PBR1 when co-expressed with AvrPphB, indicating that the individual domains and fragments of PBR1 are insufficient to trigger HR. Intriguingly, introducing the typically auto-activating D496V mutation within NB-ARC-containing fragments of PBR1 does not activate immune signaling revealing PBR1-mediated immune signaling requires cooperation of all domains *in cis*. Using co-immunoprecipitation and split-luciferase assays, we also show full-length PBR1 self-associates in the absence of AvrPphB and such self-association is not dependent on a functional P-loop/Walker A motif. Collectively, these findings provide valuable insights into PBR1-mediated disease resistance and extends upon our understanding of NLR-mediated immune signaling.

## INTRODUCTION

Plants have evolved an innate immune system that relies, in part, upon the perception of pathogen-secreted virulence (effector) proteins by intracellular immune receptors referred to as nucleotide-binding leucine-rich repeat (NLR) proteins (Jones and Dangl, 2006). Upon effector recognition, NLR proteins initiate a rapid, localized cell death response, referred to as the hypersensitive reaction (HR), that is often correlated with restricted pathogen growth and proliferation (Coll et al., 2011; Jones and Dangl, 2006; Klement and Goodman, 1967; Kourelis and van der Hoorn, 2018; Ngou et al., 2021; Ngou et al., 2022; van der Burgh and Joosten, 2019). NLR proteins conventionally have a conserved modular architecture consisting of a variable N-terminal domain, a central nucleotide-binding adaptor shared by APAF1, plant resistance proteins and CED4 (NB-ARC) domain, and a C-terminal leucine-rich repeat (LRR) domain (Caplan et al., 2008; McHale et al., 2006; Takken and Tameling, 2009).

Depending upon their N-terminal structures, NLR proteins can be further subdivided into two broad classes (Meyers et al., 2003; McHale et al., 2006). NLR proteins containing a coiled-coil (CC) domain are often referred to as CNLs (CC-NBS-LRR) while NLR proteins with a toll/interleukin-1 receptor-like (TIR) domain are known as TNLs (TIR-NBS-LRR) (Meyers et al., 1999; Cannon et al., 2002). Expression of the CC domain from several CNLs activates the hypersensitive reaction independent of effector recognition, demonstrating the CC domain plays an important role in CNL-mediated immune signaling (Bai et al., 2012; Baudin et al., 2017; Cesari et al., 2016; Collier et al., 2011; Hamel et al., 2016; Kim et al., 2018; Wang et al., 2015). Wang et al., (2019a) recently showed that the Arabidopsis disease resistance protein ZAR1 oligomerizes and forms a pinwheel-like pentamer upon activation. Upon formation of the pentameric resistosome, the N-terminal alpha helix within the CC domain of ZAR1 forms a funnel shaped structure that inserts into the plasma membrane creating a calcium-permeable channel, thereby activating a cell death response (Wang et al., 2019a). However, for some CNLs, expression of the CC domain by itself is insufficient to initiate HR-like cell death, demonstrating that immune signaling for some NLR proteins may require the cooperation of multiple domains (Ade et al., 2007; El Kasmi et al., 2017; Hamel et al., 2016). For example, the Arabidopsis CNL RPS5 does not contain N-terminal helix that is present in ZAR1 and is thus not likely to form a calcium channel (Pottinger and Innes, 2020). Instead, RPS5 contains N-terminal myristoylation and palmitoylation motifs that facilitate RPS5 localization the plasma membrane, which suggests the mechanism by which RPS5 induces immune signaling may differ from ZAR1 (Pottinger and Innes, 2020; Qi et al., 2012).

The central NB-ARC domain primarily functions in nucleotide binding, coordinating the intramolecular interactions within the NLR protein, as well as oligomerization (Collier and Moffett, 2009; Wang et al., 2019a and b). Within the NB-ARC domain, there are several conserved motifs including the P-loop/Walker A, RNBS-A (Resistance Nucleotide Binding Site-A), Kinase-2/Walker B, RNBS-B, RNBS-C, GLPL, RNBS-D, and MHD (Meyers et al., 1999). In particular, the P-loop/Walker A and MHD motifs are important for NLR function and signaling (Maekawa et al., 2011; Takken et al., 2006). Mutation of the conserved lysine residue within the P-loop/Walker A motif (GxxxxGK[T/S]) prevents ATP binding, thereby impairing NLR-mediated immune activation and the ability to confer HR-like cell death (Tameling et al., 2002; Wang et al., 2019a and b). The MHD motif is a conserved three amino acid sequence within the NB-ARC domain responsible for coordinating nucleotide binding and mutations within this motif often result in effector-independent, constitutive activation of disease resistance proteins, likely due to a reduction in ATP hydrolysis rates (Bendahmane et al., 2002; Tameling et al., 2006; Wang et al., 2019a and b).

Characterized by the repeating pattern of leucine or other hydrophobic residues, the leucine-rich repeat (LRR) domain forms an arc-shaped structure consisting of parallel beta-sheets and is often implicated in the recognition of pathogen-secreted effectors (Krasileva et al., 2010; Martin et al., 2020; Sukarta et al., 2016; Wang et al., 2019a and b). Effector recognition specificity for some NLRs can be changed by swapping the LRR domains between NLR proteins (Dodds et al., 2001; Ellis et al., 1999). Consistent with the LRR domain having a functional role in perceiving effectors, it is often the most structurally diverse region within NLR proteins and is under diversifying selection (Ellis et al., 2000; Meyers et al., 1998). In addition to effector recognition, the LRR domain has also been implicated in NLR inhibition through intramolecular interactions (Ade et al., 2007). Deletion of the LRR domain from the Arabidopsis disease resistance protein RPS5 results in constitutive activation of RPS5-dependent cell death (Ade et al., 2007). Together, these examples show that the LRR domain plays important regulatory roles through recognition and enabling conformational changes required for activation.

Carter et al., (2019) previously showed that the *Pseudomonas syringae* cysteine protease AvrPphB elicits a defense response in diverse barley varieties and this defense response is mediated by the NLR protein, *AvrPphB Response 1* (*Pbr1*). Transient co-expression of PBR1 with AvrPphB, but not an enzymatically inactive AvrPphB mutant, consistently activates cell death in *N. benthamiana*, demonstrating that PBR1 senses AvrPphB protease activity (Carter et al., 2019). In Arabidopsis, AvrPphB protease activity is recognized by RPS5 by sensing proteolytic cleavage of the serine/threonine kinase PBS1 (Ade et al., 2007; DeYoung et al., 2012; Shao et al., 2003; Zhu et al., 2004). However, PBR1 and RPS5 are not conserved orthologs and instead convergently evolved the ability to recognize AvrPphB protease activity (Carter et al., 2019). Unlike Arabidopsis RPS5, the molecular mechanisms and structural requirements leading to PBR1-dependent cell death have yet to be determined.

In this work, we characterized the induction of HR-like cell death by PBR1 in the absence and presence of the AvrPphB protease. We show that fusing an epitope tag to the N-terminus of PBR1 abolishes PBR1-mediated cell death responses. None of the individual domains or truncations of PBR1 induced a HR-like cell death response as strong as full-length PBR1 when co-expressed with AvrPphB, indicating that expression of the individual domains and fragments of PBR1 are insufficient to trigger robust HR. Furthermore, introducing the auto-activating D496V mutation into the MHD motif within NB-ARC-containing fragments of PBR1 does not activate cell death, demonstrating PBR1-mediated immune signaling requires cooperation of all domains *in cis*. Lastly, we show that PBR1 self-associates in the absence of AvrPphB and such self-association is not dependent on a functional P-loop/Walker A motif.

## MATERIALS AND METHODS

### Plant growth conditions

Seeds of *Nicotiana benthamiana* were sown in plastic pots containing ProMix Propagation Mix or Berger Seed and Propagation Mix supplemented with Osmocote slow-release fertilizer (14-14-14). Plants were maintained in a growth chamber with 16h:8h photoperiod, light:dark at 24°C with light and 20°C in the dark and 60% humidity with average light intensities at plant height of 120µmols/m^2^/s.

### Prediction of the *AvrPphB Response 1* protein structure using AlphaFold2

The predicted protein structure of *AvrPphB Response 1* (PBR1; UniProt accession A0A3Q8N4S1) was retrieved from the AlphaFold Protein Structure Database and visualized using the University of California, San Francisco (UCSF) Chimera modeler package. The PBR1 protein structure was also modeled using the Phyre2 server with the normal mode modeling setting. Phyre2 modeled residues 8-879 in PBR1 based on the template c6j5tC (Rpp13-like protein 4; ZAR1). The domain boundaries were determined using the predicted protein structure of PBR1 as determined by AlphaFold2 as well as DeepCoil, DeepCoil2, MARCOIL, and PCOILS software (Gabler et al., 2020; Zimmerman et al., 2018).

### Generation of plant expression constructs

The PBR1:sYFP, AvrPphB:myc, and AvrPphB(C98S):myc constructs have been previously described (Carter et al., 2019). Full-length PBR1, CC, CC—NB-ARC, NB-ARC, NB-ARC—LRR, and LRR fragments were PCR-amplified from the pBSDONR(P1-P4):*Pbr1.c* template (Carter et al., 2019) using either *attB*1 and *attB*4-containing primers or *attB*4r and *attB*2-containing primers. The resulting PCR products were gel-purified using the QIAquick gel extraction kit (Qiagen) and recombined into either the Gateway DONR vectors pBSDONR(P1-P4) or pBSDONR(P4r-P2) using BP Clonase II (Invitrogen) (Carter et al., 2019; Qi et al., 2012). We designated the resulting clones pBSDONR(L1-L4):*PBR1*, pBSDONR(L4r-L2):*PBR1*, pBSDONR:*CC*, pBSDONR:*NB-ARC,* pBSDONR:*LRR,* pBSDONR:*CC—NB-ARC,* and pBSDONR:*NB-ARC—LRR.* All constructs were sequence-verified to check for proper sequence and reading frame.

A commercial gene synthesis service provider (Azenta Life Sciences) was used to synthesize the following genes: PBR1 (D496V), PBR1 (K203N), CC—NB-ARC (D496V), NB-ARC (D496V), and NB-ARC—LRR (D496V). The same service provider was used to synthesize the following constructs with in-frame stop codons: PBR1, CC, CC—NB-ARC, NB-ARC, NB-ARC—LRR, and LRR fragments. The *attL*1 and *attL*4 Gateway sequences were added to the 5’ and 3’ ends, respectively, to generate Gateway-compatible DONR clones and designated pBSDONR:*PBR1* (D496V), pBSDONR:*PBR1* (K203N), pBSDONR:*CC—NB-ARC* (D496V), pBSDONR:*NB-ARC* (D496V), pBSDONR:*NB-ARC—LRR* (D496V), pBSDONR:*Pbr1.c-stop,* pBSDONR:*CC-stop*, pBSDONR:*NB-ARC-stop,* pBSDONR:*LRR-stop,* pBSDONR:*CC—NB-ARC-stop,* and pBSDONR:*NB-ARC—LRR-stop*.

To generate PBR1 protein fusions with the desired N-terminal epitope tag, pBSDONR(L4r-L2)*:PBR1* and pBSDONR(L4r-L2)*:PBR1* (D496V), was mixed with pBSDONR(L1-L4):*sYFP* (Qi et al., 2012), and the Gateway compatible, dexamethasone-inducible vector pBAV154 (Carter et al., 2019; Vinatzer et al., 2006). To generate protein fusions with the desired C-terminal epitope tags, pBSDONR:*CC*, pBSDONR:*NB-ARC,* pBSDONR:*LRR,* pBSDONR:*CC—NB-ARC,* pBSDONR:*NB-ARC—LRR*, pBSDONR:*PBR1* (D496V), pBSDONR:*NB-ARC* (D496V), pBSDONR:*CC—NB-ARC* (D496V), pBSDONR:*NB-ARC—LRR* (D496V), and pBSDONR:*PBR1* (K203N) were mixed with pBSDONR(L4r-L2):*sYFP* (Qi et al., 2012) and recombined into pBAV154. Plasmids were recombined by addition of LR Clonase II (Invitrogen) with overnight incubation at 25°C. All constructs were sequence verified and were subsequently used for transient expression assays in *Nicotiana benthamiana*.

To generate the PBR1-nLUC, PBR1-cLUC, PBR1 (K203N)-nLUC, and PBR1 (K203N)-cLUC constructs, full-length PBR1 or PBR1 (K203N) were PCR-amplified from the pBSDONR(L1-L4):*PBR1* and pBSDONR:*PBR1* (K203N) templates using *attB*1 and *attB*2-containing primers. The resulting PCR products were gel-purified using the Zymo gel extraction kit (ZymoResearch) and recombined into the Gateway DONR vector pDONR207 using BP Clonase II (Invitrogen) with overnight incubation at 25°C. The pDONR207:*PBR1* and pDONR207:*PBR1* (K203N) constructs were recombined into the Gateway-compatible pGWB-nLUC and pGWB-cLUC (Chen et al., 2008) expression vectors, which places the transgene under control of the CaMV 35S promoter. Plasmids were recombined by addition of LR Clonase II (Invitrogen) with overnight incubation at 25°C. All the constructs were verified by sequencing.

### Agrobacterium-mediated transient expression assays in *Nicotiana benthamiana*

Transient protein expression assays were performed as previously described by Carter et al., (2019) with slight modifications. Briefly, the dexamethasone-inducible constructs described above were mobilized into *Agrobacterium tumefaciens* GV3101 (pMP90) and grown on LB media plates containing 25µg of gentamicin sulfate and 50µg of kanamycin per milliliter for 2 days at 30°C. Cultures were prepared in liquid media (10 ml) supplemented with the appropriate antibiotics and were shaken overnight at 30°C at 225rpm on an orbital shaker. Following overnight incubation, the cells were pelleted by centrifuging at 3,700rpm for 3 min and resuspended in 10mM MgCl_2_, adjusted to an OD_600_ of 0.3 for cell death and electrolyte leakage assays and an OD_600_ of 0.1 for immunoblot assays (final concentration for each strain in a mixture) and were incubated with 100µM acetosyringone (Sigma-Aldrich) for 2-4 hours at room temperature. For co-expression of multiple constructs, the suspensions were mixed in equal ratios and infiltrated into the underside of 3-week-old *Nicotiana benthamiana* leaves with a needleless syringe. Tissue samples for protein extraction were harvested 6 hours after dexamethasone application, flash frozen in liquid nitrogen and stored at −80°C. Leaves for macroscopic cell death were evaluated 24 hours following transgene induction and were removed for photography shortly thereafter. Unless otherwise noted, all experiments were repeated at least three independent times.

### Electrolyte leakage assay in *N. benthamiana*

To quantify cell death, four leaf discs were collected from *N. benthamiana* leaves using a cork borer (5mm diameter) 2 hours post-dexamethasone application. Four leaf discs collected from individual leaves of four different plants were included for each experimental replicate. The leaf discs were briefly washed with deionized water and floated in 5ml of deionized water supplemented with 0.001% Tween 20 (Sigma-Aldrich) in 12-well tissue culture plates. Electrolyte leakage was monitored using a Traceable Pen conductivity meter (VWR) at the indicated time points following dexamethasone application.

Statistical analyses were performed using the conductivity data from the last time point collected. One-way ANOVA followed by Tukey’s honestly significant difference (HSD) tests (using α = 0.05) were performed to identify significant differences between the measurements. Measurements with no statistically significant differences are indicated with the same letter whereas different letters represent significant differences.

### Immunoblot analyses

Frozen leaf tissue was ground in two volumes of protein extraction buffer (150 mM NaCl, 50 mM Tris [pH 7.5], 0.1% Nonidet P-40 [Sigma-Aldrich], 1% plant protease inhibitor cocktail [Sigma-Aldrich], and 1% 2,2’-dipyridyl disulfide [Chem-Impex]) using a cold ceramic mortar and pestle. The cell debris was pelleted at 10,000xg for 12 min at 4°C. Seventy-five microliters of the collected supernatants were combined with 25 microliters of 4x Laemmli buffer (BioRad) supplemented with 10% β-mercaptoethanol and the mixture was boiled at 95°C for 10 minutes. All samples were separated on a 4-20% Tris-glycine stain free polyacrylamide gel (BioRad) at 180V for 1 hour in 1x Tris/glycine/SDS running buffer. Total proteins were transferred to a nitrocellulose membrane (GE Water and Process Technologies). Ponceau staining was used to confirm equal loading and transfer of total protein samples. Membranes were washed with 1x Tris-buffered saline (50mM Tris-HCl, 150mM NaCl, pH 7.5) solution supplemented with 0.1% Tween20 (TBST) and blocked in 5% skim milk (Becton, Dickinson & Company) for at least 1 hour at room temperature. Proteins were subsequently detected with either horseradish peroxidase (HRP)-conjugated anti-c-myc antibody (1:5,000) (Thermo Fisher Scientific) or HRP-conjugated anti-GFP antibody (1:5,000) (Miltenyi Biotec). Membranes were washed three times for 15 minutes in 1x TBST and immunoblots were incubated for 5 minutes at room temperature with either Clarity Western ECL (BioRad) or Supersignal West Femto maximum sensitivity substrates (Thermo Scientific). Immunoblots were developed using either X-ray film or an ImageQuant 500 CCD imaging system (Cytiva).

### Co-immunoprecipitation analyses

Co-immunoprecipitation assays were performed as previously described by Carter et al., (2019) with slight modifications. Briefly, *N. benthamiana* leaf tissue transiently expressing either sYFP or myc-tagged proteins were ground in 1mL of IP buffer (150 mM NaCl, 50 mM Tris [pH 7.5], 10% glycerol, 1 mM dithiothreitol, 1 mM EDTA, 1% Nonidet P-40, 0.1% Triton X-100, 1% plant protease inhibitor cocktail, and 1% 2,2’-dipyridyl disulfide) using a cold ceramic mortar and pestle. Homogenates were centrifuged at 12,000 x g for 12 minutes at 4°C twice to remove plant cell debris. Five hundred microliters of the supernatant was incubated with 10µL of green fluorescent protein (GFP)-Trap A (Chromotek) bead slurry for 3 hours at 4°C with constant end-over-end rotation. Following the 3-hour incubation, the bead slurry was pelleted by centrifuging at 1,000 x g for 1 minute at 4°C and washed 5 times with 500µL of the IP wash buffer. The immunoprecipitates were resuspended in IP buffer and combined with 4x Laemmli buffer (BioRad) supplemented with 10% β-mercaptoethanol and the mixtures were boiled at 95°C for 5 min. Protein samples were separated on 4-20% Tris-glycine stain-free polyacrylamide gels (Bio-Rad) at 180 V for 1 hour in 1X Tris/glycine/SDS running buffer. Total proteins were transferred to nitrocellulose membranes (GE Water and Process Technologies) at 100 V for one hour. Membranes were washed with 1X Tris-buffered saline (50 mM Tris-HCl, 150 mM NaCl [pH 6.8]) solution containing 0.1% Tween20 (TBST) and incubated with 5% Difco skim milk for 1 hour at room temperature with gentle shaking. Proteins were subsequently detected with horseradish peroxidase (HRP)-conjugated anti-GFP antibody (1:5,000) (Miltenyi Biotec) or HRP-conjugated anti-c-myc antibody (1:5,000) (Thermo Fisher Scientific) for 1 hour at room temperature with gentle shaking. Following antibody incubation, membranes were washed at least three times for 15 minutes in 1x TBST solution and incubated for 5 minutes at room temperature with either Clarity Western ECL (BioRad) or Supersignal West Femto maximum sensitivity substrates (Thermo Scientific). Immunoblots were developed using either X-ray film or an Amersham ImageQuant 500 CCD imaging system (Cytiva).

### Split-luciferase (split-LUC) assays in *N. benthamiana*

The split-luciferase complementation imaging assay (LCI) assay was performed as previously described (Chen et al., 2008) with modifications. *A. tumefaciens* harboring the nLUC- and cLUC-tagged fusion constructs were agroinfiltrated into *N. benthamiana* and split-luciferase assays were performed 2 days following agroinfiltration. For CCD imaging, *N. benthamiana* leaves were sprayed with 1mM luciferin dissolved in deionized water and subsequently kept in the dark for 10 minutes to quench detection of background luciferase activity. A low-light cooled CCD imaging system (CHEMIPROHT 1300B/LND, 16 bits; Roper Scientific) was used to capture the luciferase activity using an exposure time of 1 minute with 3×3 binning.

To quantify the luciferase signal, 6-10 leaf discs (5 mm diameter) were collected two days following agroinfiltration and transferred to a 24-well microtiter plate containing deionized water supplemented with 1mM luciferin. The microtiter plates were wrapped in aluminum foil for 10 minutes to remove background luminescence signals. Luciferase activity was recorded using an Infinite M200 Pro Plate Reader (Tecan) using luminescence detection parameters for at least 40 minutes. Each data point consisted of three replicates and three independent experiments were performed.

## RESULTS

### *In silico* PBR1 protein structure prediction

The *AvrPphB Response 1* (PBR1) protein sequence encodes a full-length NLR protein consisting of 939 amino acids with an N-terminal coiled-coil (CC) domain (amino acids 1-144), a central nucleotide-binding (NB-ARC) domain (amino acids 145-508), and a C-terminal leucine-rich repeat (LRR) domain (amino acids 509-939) consisting of 13 repeats (Figure 1A). The conserved EDVID motif (EDCMD in PBR1) can be identified within the CC domain of PBR1 along with all expected motifs that define the NB-ARC domain (Figure 1A) (Meyers et al., 1999). We retrieved the predicted protein structure of PBR1 from the AlphaFold Protein Structure Database to better understand the structural dynamics of PBR1. Visual inspection shows that it adopts a compact modular structure with the N-terminal coiled-coil domain forming a four-helical barrel conformation similar to other CNLs (Figure 1B; Casey et al., 2016; Maekawa et al., 2011; Wang et al., 2019a and b). Furthermore, AlphaFold2 confidently predicts the conformation and positioning of the NB-ARC and LRR domains relative to the coiled-coil domain (Figure 1B). Intriguingly, the most C-terminal portion of PBR1 (amino acids 922-939) is predicted to be unstructured and contains a non-LRR extension (Figure 1B).

**Figure 1.**
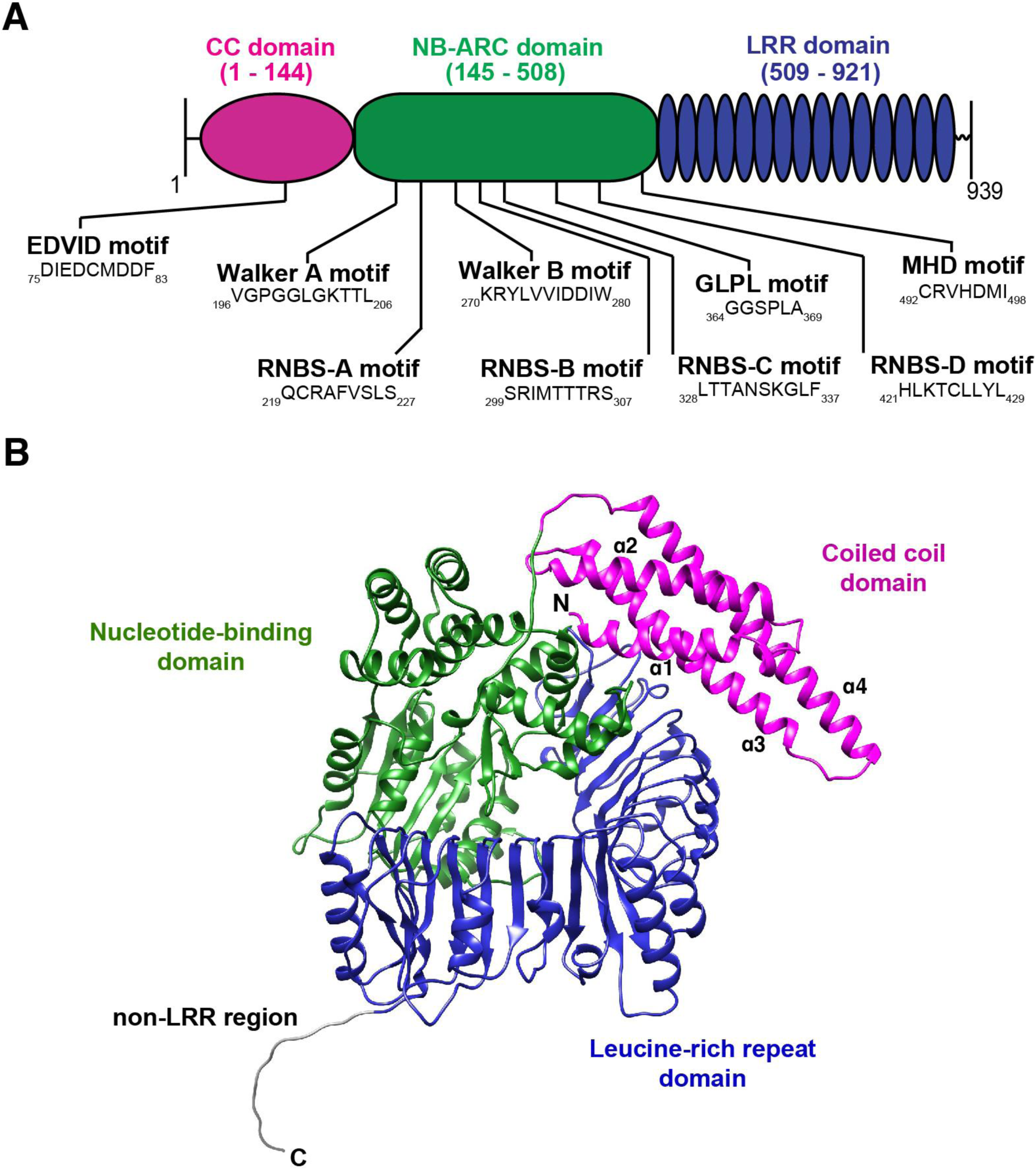
Schematic illustration and predicted protein structure of barley PBR1. **(A)** The predicted coiled-coil (CC; amino acids 1-144), nucleotide-binding (NB-ARC; amino acids 145-508), and leucine-rich repeat (LRR; amino acids 509-883) domains of PBR1 are indicated by magenta, green, and blue bars, respectively. The conserved motifs within the NB-ARC domain (Walker A, RNBS-A, Walker B, RNBS-B, GLPL, RNBS-D, and MHD) were predicted using NLRexpress (https://nlrexpress.biochim.ro) (Martin et al., 2022) and shown beneath the schematic illustration. Numbers indicate amino acid positions. **(B)** The predicted protein structure of PBR1 as determined by AlphaFold2. The predicted protein structure of PBR1 (UniProt accession A0A3Q8N4S1) was retrieved from the AlphaFold Protein Structure Database and visualized using the University of California, San Francisco (UCSF) Chimera modeler package. The predicted coiled-coil, nucleotide-binding, and leucine-rich repeat domains are color-coded magenta, green, and blue, respectively. The non-LRR extension (amino acids 922-939) is shown in gray.

### An N-terminal epitope tag interferes with PBR1-mediated immune signaling

We previously showed that fusing the super yellow fluorescent protein (sYFP) epitope tag to the C-terminus of PBR1 (PBR1:sYFP) confers a strong cell death response in *N. benthamiana* when co-expressed with myc-tagged AvrPphB (Carter et al., 2019). To further dissect defense signaling initiated by the PBR1 immune receptor, we fused the sYFP protein to the N-terminus of PBR1 (sYFP:PBR1) and coexpressed with AvrPphB:myc in *N. benthamiana* (Figure 2A). Consistent with the observations of Carter et al., (2019), our positive control of transiently coexpressing PBR1:sYFP with AvrPphB:myc resulted in a robust cell death response. However, expression of sYFP:PBR1 with AvrPphB:myc failed to consistently induce observable tissue collapse 24 hours post-transgene induction (Figure 2B). We next performed electrolyte leakage analyses to better quantify the cell death responses. Consistent with the macroscopic symptoms, coexpression of PBR1:sYFP with AvrPphB:myc resulted in significant ion leakage whereas coexpression of sYFP:PBR1 with AvrPphB:myc failed to induce significant ion leakage in *N. benthamiana* (Figure 2C). In addition, expression of sYFP:PBR1 by itself failed to produce significant electrolyte leakage as observed with PBR1:sYFP, further demonstrating that N-terminally tagged PBR1 is not capable of activating a cell death response. To rule out the possibility that the lack of observable cell death was due to insufficient protein accumulation of sYFP:PBR1, we performed an immunoblot analysis to assess protein accumulation and integrity (Figure 2D). PBR1:sYFP coexpressed with AvrPphB:myc does not produce detectable protein accumulation, likely a result of rapid protein degradation from activation of the HR. In contrast, we detected protein accumulation of both sYFP:PBR1 and AvrPphB:myc by immunoblot blot analyses 6 hours post-transgene induction, demonstrating that the absence of cell death with these constructs is not due to a lack of protein accumulation.

**Figure 2.**
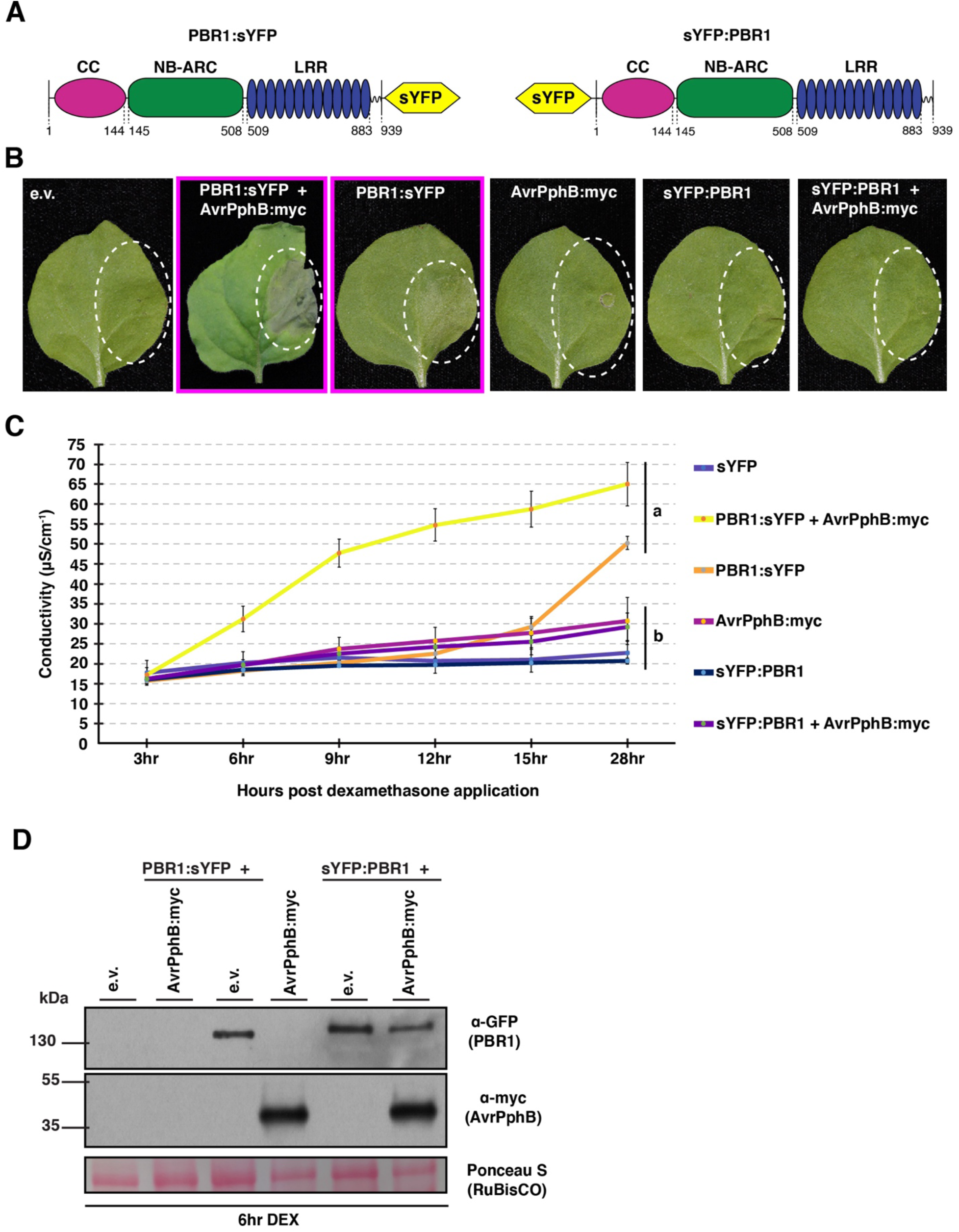
N-terminal tagged PBR1 fails to activate HR-like cell death in response to AvrPphB protease activity in *N. benthamiana*. **(A)** Schematic illustration of the N-terminal and C-terminal sYFP-tagged PBR1 constructs used in panels B-D. The coiled-coil (CC) domain is shown in magenta, the nucleotide-binding (NB-ARC) domain is shown in green, and the leucine-rich repeat (LRR) domain is shown in blue. The super yellow fluorescent protein (sYFP) is shown in yellow. Numbers located beneath the schematic delineate amino acid positions. **(B)** Fusing an epitope tag to the N-terminus of full-length PBR1 abolishes cell death-inducing activity when transiently co-expressed with AvrPphB in *N. benthamiana*. The indicated construct combinations were infiltrated into 3-week-old *N. benthamiana* using *Agrobacterium tumefaciens*-mediated transformation (agroinfiltration). C-terminal-tagged PBR1 co-expressed with myc-tagged AvrPphB was used as a positive control (Carter et al., 2019). White circles indicate the agroinfiltration areas within the *N. benthamiana* leaves. Phenotypes were observed 24 hours after dexamethasone induction and a representative leaf was photographed under white light shortly thereafter. Photographs outlined with a magenta box indicate the agroinfiltrated leaf area showed observable cell death. **(C)** Plant cell death symptoms were quantified using electrolyte leakage (ion leakage/conductivity). Agroinfiltration was used to transiently express the indicated construct combinations in *N. benthamiana*. C-terminal tagged PBR1 co-expressed with myc-tagged AvrPphB was used as a positive control (Carter et al., 2019). Conductivity was recorded at the indicated timepoints following transgene induction and is shown as mean SE (n = 4). Statistical significance was assessed using one-way ANOVA followed by Tukey-Kramer honestly significant difference analyses. Different letters represent significantly different means based on Tukey-Kramer honestly significant difference test (using α = 0.05). Statistical analyses were performed using the conductivity (µS/cm^-1^) data from the last timepoint. **(D)** Detectable protein expression of N-terminal and C-terminal sYFP-tagged PBR1 constructs. The indicated constructs were transiently expressed in 3-week-old *N. benthamiana* using agroinfiltration. All proteins were under the control of a dexamethasone inducible promoter. Total protein was extracted 6 hours following dexamethasone application. Equal amounts of total protein were resolved on 4-20% SDS-PAGE gels, blotted onto nitrocellulose membranes, and probed using either anti-GFP-horseradish peroxidase or anti-Myc-horseradish peroxidase antibodies. Ponceau staining of the RuBisCO large subunit was used as a loading control.

Carter et al., (2019) showed that PBR1 mediates indirect recognition of AvrPphB protease activity likely by sensing cleavage of an endogenous *N. benthamiana* PBS1 or PBS1-like (PBL) protein. Hence, fusing an epitope tag to the N-terminus of PBR1 may interfere with its ability to detect the cleaved PBS1 or PBS1-like protein, thus inhibiting PBR1-mediated cell death. To interrogate this hypothesis, we constructed an auto-active allele of PBR1 wherein the aspartic acid residue within the MHD motif was replaced with valine, generating PBR1 (D496V) (Supplemental Figure 1A). Mutation of the conserved MHD motif to MHV often results in constitutive activation of NLR proteins (Bendahmane et al., 2002; Tameling et al., 2006; Wang et al., 2019a and b). Transient expression of this allele in *N. benthamiana* confirmed that PBR1 (D496V) is indeed auto-active as indicated by the macroscopic cell death 24 hours post-transgene induction (Supplemental Figure 1B). Following fusion to sYFP, we observed that expression of PBR1 (D496V):sYFP, but not sYFP:PBR1 (D496V), induced a robust cell death response in *N. benthamiana* (Supplemental Figure 1C). Immunoblot analyses also confirmed protein expression of the sYFP:PBR1 (D496V) derivative, thus the lack of cell death is likely not a result of insufficient protein accumulation (Supplemental Figure 1D). Collectively, these data demonstrate that an N-terminal epitope tag disrupts PBR1-mediated immune signaling.

### Immune signaling requires full-length PBR1

The interference of an N-terminal tag with PBR1-mediated immune signaling led us to hypothesize the N-terminus may have a functional role in inducing cell death. Indeed, the CC domain of several NLR proteins has been shown to induce HR-like cell death when transiently expressed, indicating their importance in CNL-mediated immune signaling (Bai et al., 2012; Baudin et al., 2017; Cesari et al., 2016; Collier et al., 2011; Hamel et al., 2016; Kim et al., 2018; Wang et al., 2015). To better understand the involvement of PBR1 domains in initiating cell death in *N. benthamiana*, we generated constructs overexpressing PBR1 fragments with C-terminal sYFP tags, including individual protein domains, based on the predicted secondary structure of PBR1: CC (amino acids 1-144), NB-ARC (amino acids 145-508), and LRR (amino acids 509-939) domains as well as the CC—NB-ARC (amino acids 1-508) and NB-ARC—LRR (amino acids 145-939) fragments (Figure 3A). Upon transient expression of the sYFP fusion proteins with either free sYFP or myc-tagged AvrPphB in *N. benthamiana*, we consistently observed that none of the individual domains or fragments of PBR1 induced HR-like cell death equivalent to that of full length PBR1:sYFP coexpressed with AvrPphB:myc (Figure 3B; Supplemental Figure 2A). Electrolyte leakage analyses confirmed the lack of cell death induction and all constructs accumulated detectable protein 6 hours post-transgene induction (Figure 3C-D; Supplemental Figure 2B-C). Overexpression of non-epitope tagged individual domains or truncations with or without AvrPphB also did not lead to the induction of observable cell death, indicating that the fused sYFP was likely not inhibiting functionality (Supplemental Figure 3). Taken together, these data suggest the individual domains or fragments of PBR1 are unable to activate a cell death response when coexpressed with or without AvrPphB, regardless of the presence of an epitope tag.

**Figure 3.**
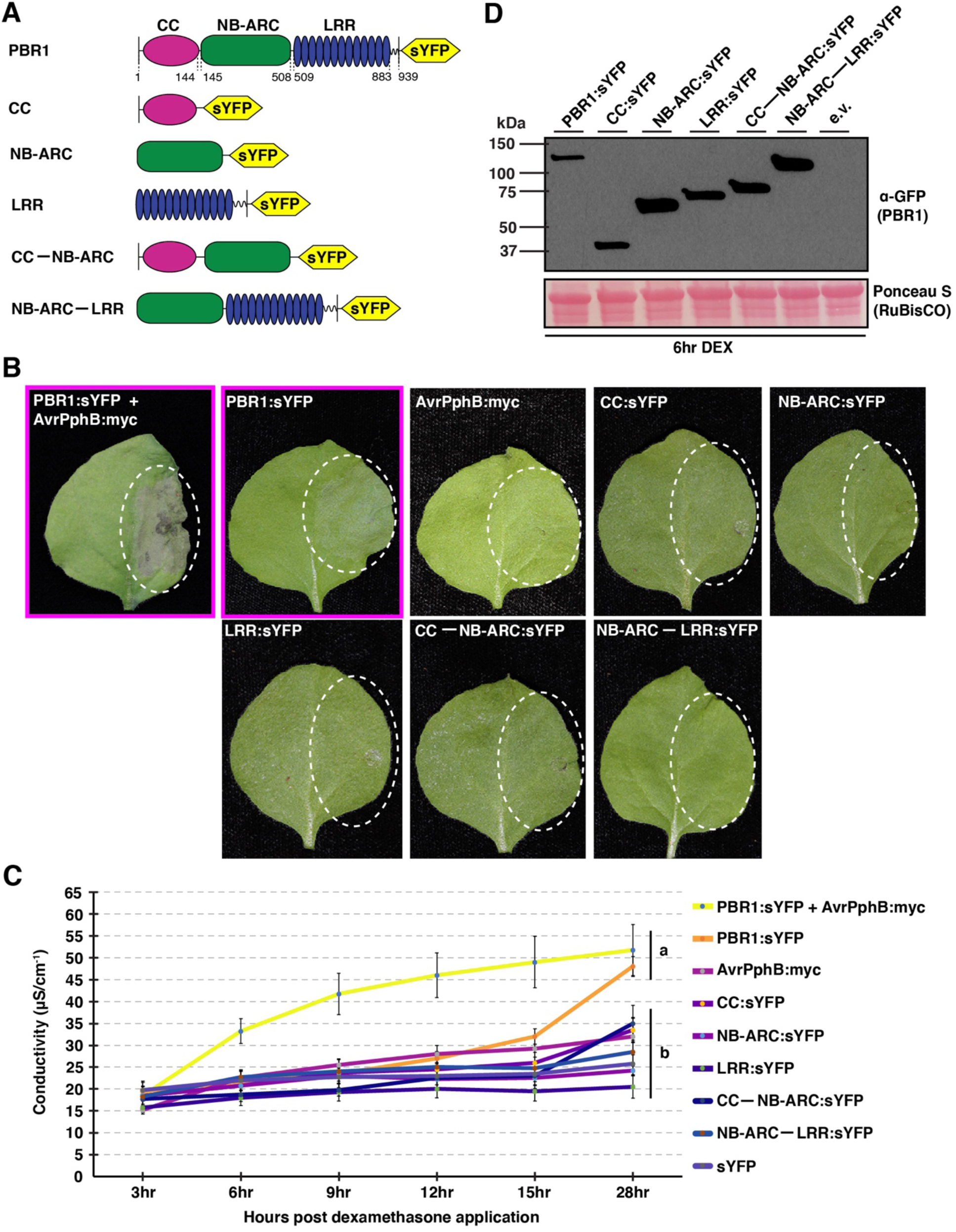
Truncation variants of PBR1 fail to induce HR-like cell death in *N. benthamiana*. **(A)** Schematic representation of full-length PBR1 and PBR1 domains and truncations used throughout this work. The coiled-coil (CC), nucleotide binding (NB-ARC), and leucine-rich repeat (LRR) domains were determined using secondary structure predictions and amino acid sequence comparisons to other plant NLR proteins. Numbers beneath the schematic illustration are the amino acid locations delineating the boundaries of the CC, NB-ARC, and LRR domains. PBR1 domains and fragments are shown below the full-length PBR1 schematic. **(B)** Lack of observable cell death induction by sYFP-tagged PBR1 domains and fragments when transiently expressed in *N. benthamiana*. Agroinfiltration was used to transiently co-express the indicated constructs into 3-week-old *N. benthamiana*. All proteins were C-terminally tagged with sYFP and under the control of a dexamethasone inducible promoter. PBR1:sYFP co-expressed with AvrPphB:myc was used as a positive control. White circles indicate the agroinfiltration areas within the *N. benthamiana* leaves. Leaves were harvested 24 hours post-transgene induction and a representative leaf was photographed under white light shortly thereafter. **(C)** Plant cell death was quantified using electrolyte leakage. Agroinfiltration was used to transiently express the indicated construct combinations in *N. benthamiana*. Conductivity was recorded at the indicated timepoints following transgene induction and is shown as mean SE (n = 4). Statistical significance was assessed using one-way ANOVA followed by Tukey-Kramer honestly significant difference analyses. Different letters represent significantly different means based on Tukey-Kramer honestly significant difference test (using α = 0.05). Statistical analyses were performed using the conductivity (µS/cm^-1^) data from the last timepoint. **(D)** Protein accumulation of sYFP-tagged PBR1 domains and truncation variants as well as full-length PBR1 from experiment shown in panel (A). The indicated constructs were transiently expressed in 3-week-old *N. benthamiana* using agroinfiltration, and total protein was extracted 6 hours following transgene induction and analyzed by immunoblotting using an anti-GFP-horseradish peroxidase antibody. Ponceau staining of the RuBisCO large subunit was used as a loading control. Expected M_r_s of proteins: PBR1:sYFP ∼133 kDa, CC:sYFP ∼43kDa, NB-ARC:sYFP ∼67kDa, LRR:sYFP ∼76kDa, CC—NB-ARC:sYFP ∼85kDa, and NB-ARC—LRR:sYFP ∼116kDa.

Transient coexpression of complementary domains (i.e. CC—NB-ARC fragment with the LRR domain) *in trans* can often reconstitute a functional disease resistance protein capable of immune signaling, as was observed with the disease resistance proteins Rx and Mi-1.2 from potato and tomato, respectively (Moffett et al., 2002; van Ooijen et al., 2008). We, therefore, investigated whether coexpression of non-epitope tagged complementary domains or fragments *in trans* would activate cell death. Unlike Rx and Mi-1.2, transient coexpression of individual domains or fragments *in trans* with or without AvrPphB failed to activate macroscopic cell death 24 hours post-transgene induction (Supplemental Figure 4). Collectively, our results indicate PBR1-mediated immune signaling cannot be recapitulated by the *in trans* expression of the individual domains or truncations and requires cooperation of all domains.

### Introducing the auto-activating D496V mutation within the NB-ARC domain-containing fragments of PBR1 does not induce HR-like cell death

Having shown PBR1 (D496V) elicits an effector-independent cell death response (Supplemental Figure 1), we next investigated whether introducing this mutation within the NB-ARC domain-containing fragments would render these truncations auto-active as previously observed with the wheat CNL Sr35 (Bolus et al., 2020) (Figure 4A). The NB-ARC (D496V):sYFP, CC—NB-ARC (D496V):sYFP, and NB-ARC—LRR (D496V):sYFP derivatives were unable to activate a cell death response equivalent to that of PBR1:sYFP (D496V) or PBR1:sYFP coexpressed with AvrPphB:myc (Figure 4B). Likewise, electrolyte leakage analyses confirmed the lack of cell death induction by the NB-ARC (D496V):sYFP, CC—NB-ARC (D496V):sYFP, and NB-ARC—LRR (D496V):sYFP derivatives (Figure 4C).

**Figure 4.**
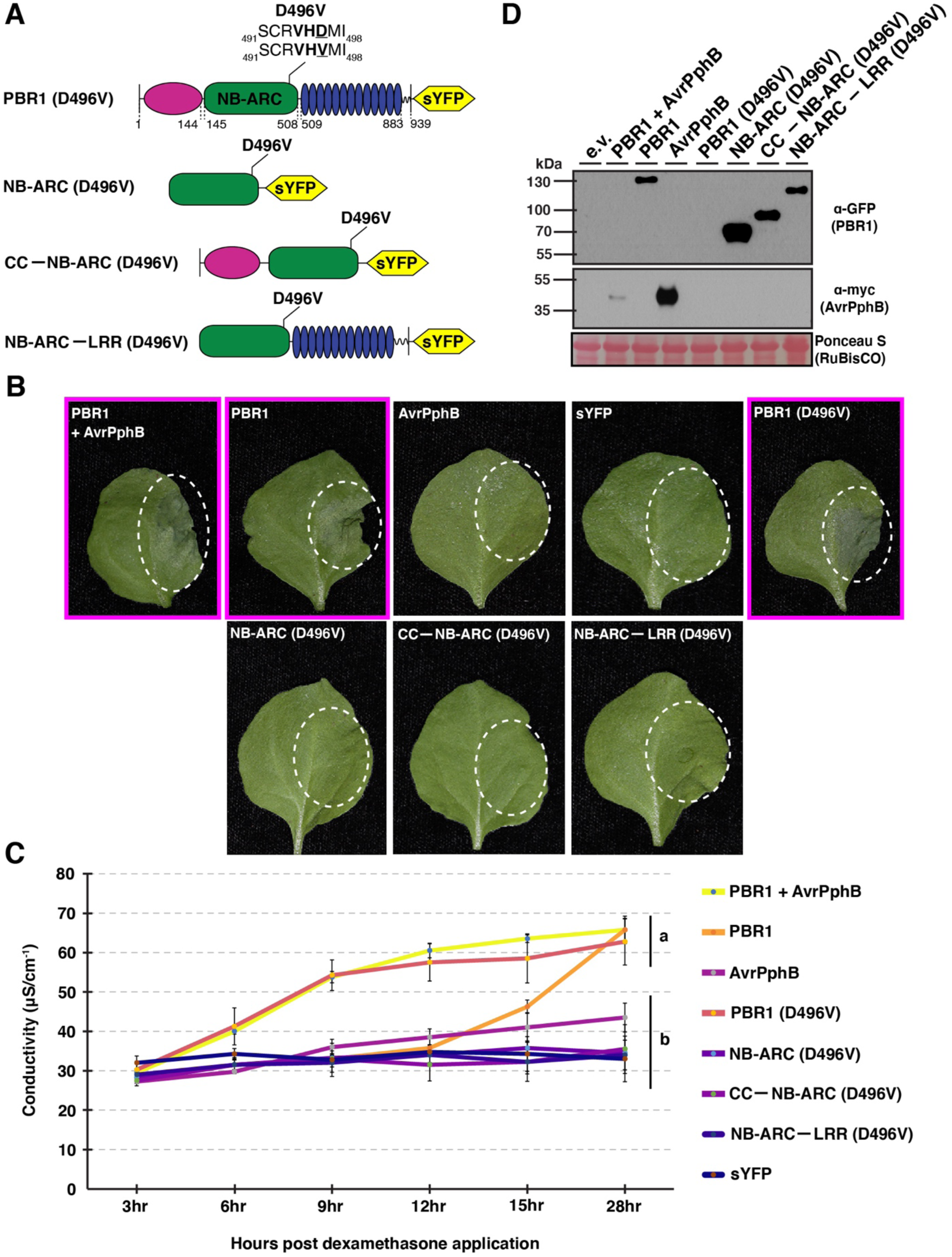
Introducing the D496V mutation within the sYFP-tagged NB-ARC, CC— NB-ARC, and NB-ARC—LRR fragments fails to active cell death in *N. benthamiana*. **(A)** Schematic representation of the sYFP-tagged full-length PBR1 (D496V), NB-ARC (D496V), CC—NB-ARC (D496V), and NB-ARC—LRR (D496V) derivatives used in panels B-D. The MHD motif is shown in bold and the D496V mutation is underlined. All proteins were C-terminally tagged with sYFP and under the control of a dexamethasone inducible promoter. **(B)** The PBR1 (D496V) derivative induces macroscopic cell death whereas the NB-ARC (D496V) domain and CC—NB-ARC (D496V) and NB-ARC—LRR (D496V) fragments fail to induce HR-like cell death when transiently expressed in *N. benthamiana*. Agroinfiltration was used to transiently co-express the indicated constructs into 3-week-old *N. benthamiana*. Full-length PBR1 co-expressed with AvrPphB was used as a positive control. White circles indicate the agroinfiltration areas within the *N. benthamiana* leaves. Leaves were harvested 24 hours post-transgene induction and a representative leaf was photographed under white light shortly thereafter. Photographs outlined with a magenta box indicate the agroinfiltrated leaf area showed observable cell death. **(C)** Plant cell death symptoms were quantified using electrolyte leakage from duplicates of the leaves shown in panel (C). Conductivity was recorded at the indicated timepoints following transgene induction and is shown as mean SE (n = 4). Statistical significance was assessed using one-way ANOVA followed by Tukey-Kramer honestly significant difference analyses. Different letters represent significantly different means based on Tukey-Kramer honestly significant difference test (using α = 0.05). Statistical analyses were performed using the conductivity (µS/cm^-1^) data from the last timepoint. **(D)** Protein accumulation of the sYFP-tagged the NB-ARC (D496V), CC—NB-ARC (D496V), and NB-ARC—LRR (D496V) derivatives. The indicated constructs were transiently expressed in 3-week-old *N. benthamiana* using agroinfiltration. Total protein was extracted 6 hours following dexamethasone application. Equal amounts of total protein were resolved on 4-20% SDS-PAGE gels, blotted onto nitrocellulose membranes, and probed using either anti-GFP-horseradish peroxidase or anti-Myc-horseradish peroxidase antibodies. Ponceau staining of the RuBisCO large subunit was used as a loading control.

To rule out problems with protein stability of the sYFP fusion proteins, we performed anti-GFP and anti-myc immunoblot analyses. The PBR1 (D496V):sYFP derivative failed to accumulate detectable protein 6 hours post-transgene induction, likely a result of rapid HR-like cell death (Figure 4D). Importantly, the NB-ARC (D496V):sYFP, CC—NB-ARC (D496V):sYFP, and NB-ARC—LRR (D496V):sYFP constructs accumulated detectable protein, demonstrating that the lack of cell death with the sYFP-tagged constructs is not due to a lack of sufficient protein accumulation (Figure 4D). Our findings further support the observation that the individual PBR1 domains are unable to activate immune signaling.

### Full-length PBR1 self-associates in the absence of AvrPphB

It is well known that self-association (oligomerization) of NLR proteins is essential for NLR-mediated immune signaling (Krasileva et al., 2010; Wang, et al., 2015). To investigate whether full-length PBR1 self-associates in the absence of AvrPphB, we fused the 5xmyc epitope tag to the C-terminus of PBR1 (PBR1:myc), co-expressed with PBR1:sYFP, and performed co-immunoprecipitation (co-IP) assays in *N. benthamiana.* As a control, we co-expressed free sYFP with PBR1:myc. In the absence of AvrPphB, PBR1:sYFP and PBR1:myc consistently co-immunoprecipitated (Figure 5A). To further validate the *in planta* self-association of PBR1, we performed split-luciferase complementation (Split-LUC) assays. Specifically, we fused the N-terminal half of luciferase (nLUC) and C-terminal half of luciferase (cLUC) to full-length PBR1, then transiently co-expressed PBR1-nLUC with PBR1-cLUC in *N. benthamiana*. Consistent with the co-immunoprecipitation results, co-expression of PBR1-nLUC with PBR1-cLUC resulted in detectable luciferase signal, whereas nLUC co-expressed with cLUC resulted in weaker luciferase expression (Figure 5B and 4C). Taken together, these data indicate PBR1 self-associates in the absence of AvrPphB inside plant cells.

**Figure 5.**
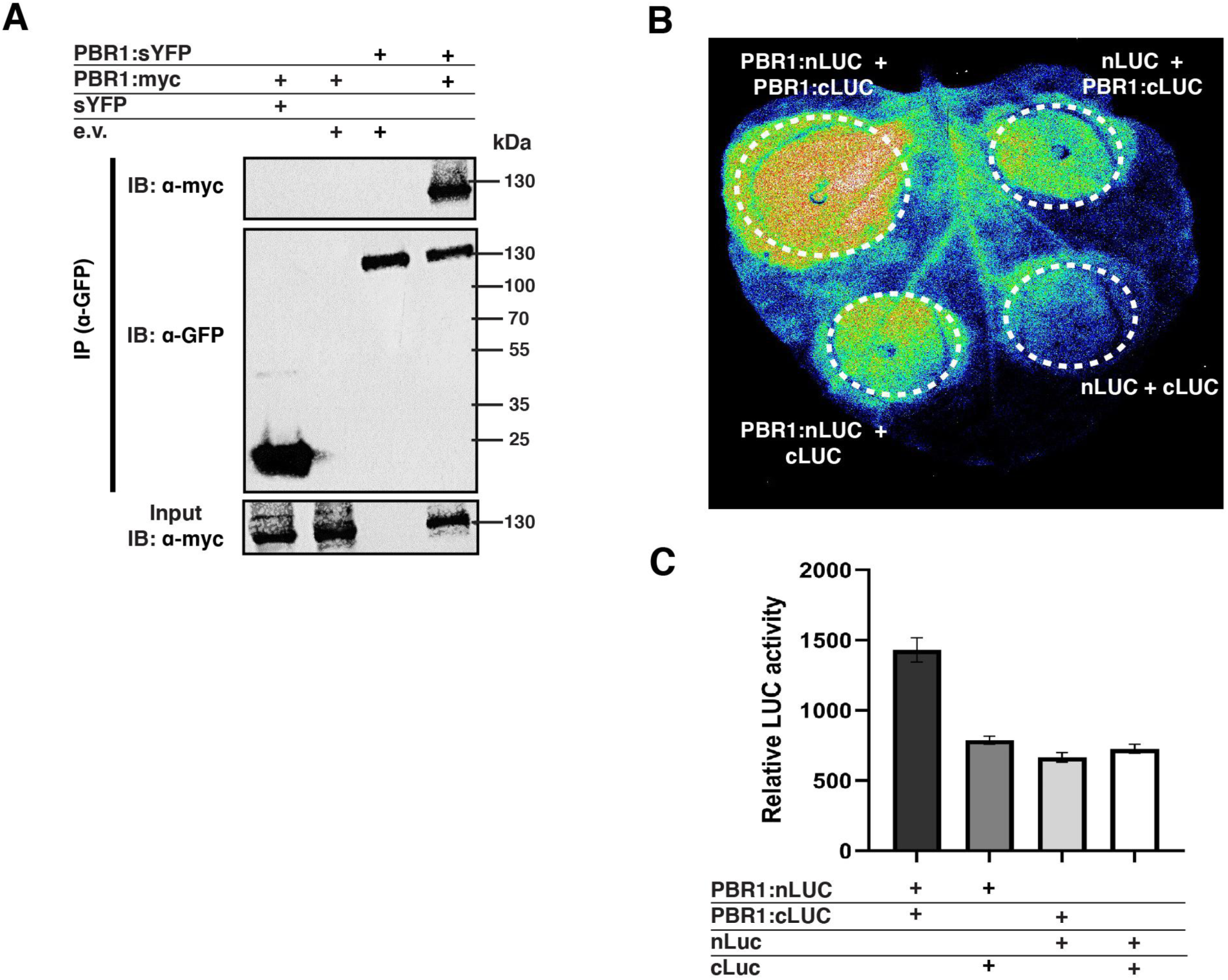
Full-length PBR1 self-associates in the absence of AvrPphB in *N. benthamiana*. **(A)** Full-length PBR1 self-associates prior to activation by AvrPphB. The indicated constructs were transiently expressed in *N. benthamiana* using agroinfiltration. Total protein was extracted 6 hours following dexamethasone application. The indicated protein samples were immunoprecipitated (IP) by Green Fluorescent Protein (GFP)- Trap agarose bead slurry, followed by immunoblotting (IB) with the indicated antibodies. **(B)** PBR1 self-associates in a firefly luciferase complementation imaging assay. PBR1:nLUC and PBR1:cLUC were transiently coexpressed in *N. benthamiana* using agroinfiltration. Thirty hours following agroinfiltration, *N. benthamiana* leaves were sprayed with 1mM luciferin and a low-light cooled CCD imaging system was used to capture luciferase activity. Free cLUC coexpressed with PBR1:nLUC, free nLUC coexpressed with PBR1:cLUC, and free nLUC coexpressed with free cLUC were used as negative controls. **(C)** Relative luciferase activity was quantified using a microplate luminescence reader (Infinite M200 Pro Plate Reader). Transient expression of the indicated constructs and luciferase activity was detected as described in panel (B). Each data point consisted of three replicates and three independent experiments were performed.

### Inter- and intra-molecular associations of PBR1 domains and truncations

Having shown full-length PBR1 self-associates, we next investigated whether the isolated domains and truncations of PBR1 could also self-associate. Indeed, the coiled-coil domain from several NLR proteins has been shown to form homodimers (Ade et al., 2007; Baudin et al., 2017; Casey et al., 2016; Cesari et al., 2016; El Kasmi et al., 2017; Wang et al., 2015). To test our hypothesis, we generated myc-tagged PBR1 domains and transiently co-expressed them with their corresponding sYFP-tagged PBR1 domains in *N. benthamiana*. Consistent with our hypothesis, the co-immunoprecipitation experiments revealed that each domain or fragment physically associates with itself in the absence of AvrPphB (Figure 6A). We also assessed which PBR1 domains associated with each other using co-immunoprecipitation assays. These co-IP experiments revealed the CC:sYFP associated with the NB-ARC:myc and LRR:myc, and the NB-ARC:sYFP domain interacted with the LRR:myc (Supplemental Figure 5).

**Figure 6.**
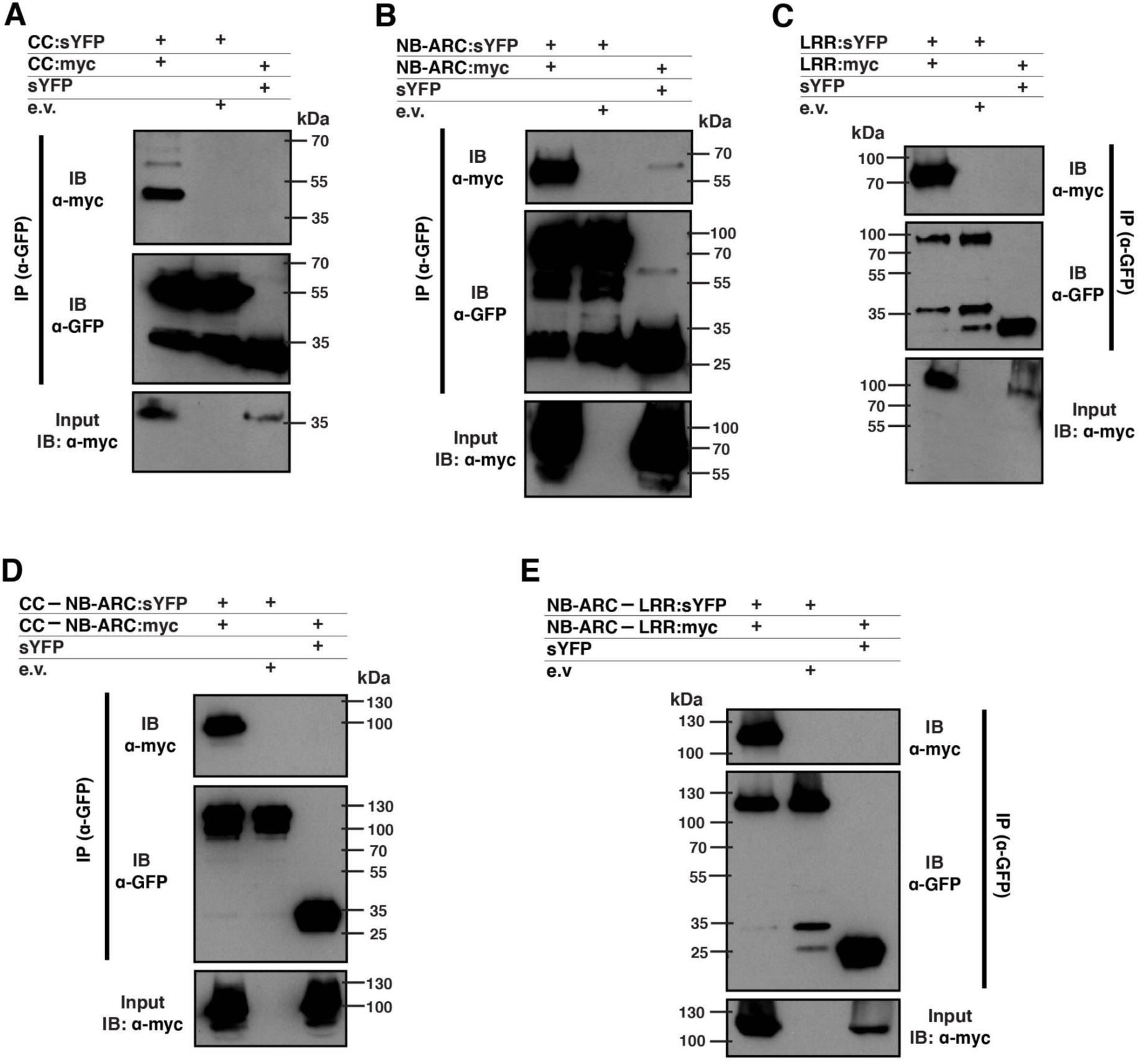
Self-association of the PBR1 domains and truncations. Co-immunoprecipitation experiments were performed to detect the self-association of the CC, NB-ARC, LRR, CC—NB-ARC, and NB-ARC—LRR fragments. The indicated constructs were transiently expressed in *N. benthamiana* and total protein was extracted 6 hours post-transgene induction. Protein samples were immunoprecipitated by GFP-Trap agarose bead slurry, followed by immunoblotting with the indicated antibodies.

### Introducing the K203N mutation abolishes PBR1-mediated cell death but not self-association of full-length PBR1

To further investigate PBR1 self-association, we next tested whether such homodimerization was dependent upon a functional P-loop/Walker A motif in the NB-ARC domain. In contrast to the autoactive MHD motif mutant, mutating the conserved lysine residue within the P-loop/Walker A motif abolishes nucleotide binding, thereby inhibiting NLR-mediated activation of cell death (Takken et al., 2006). Prior to testing our hypothesis, we mutated the lysine residue within the P-loop/Walker A motif to asparagine, thereby generating PBR1 (K203N). Transient expression of this allele in *N. benthamiana* confirmed that PBR1 (K203N) fails to activate cell death when co-expressed with AvrPphB in *N. benthamiana*, despite expressing stably (Figure 7A-C).

**Figure 7.**
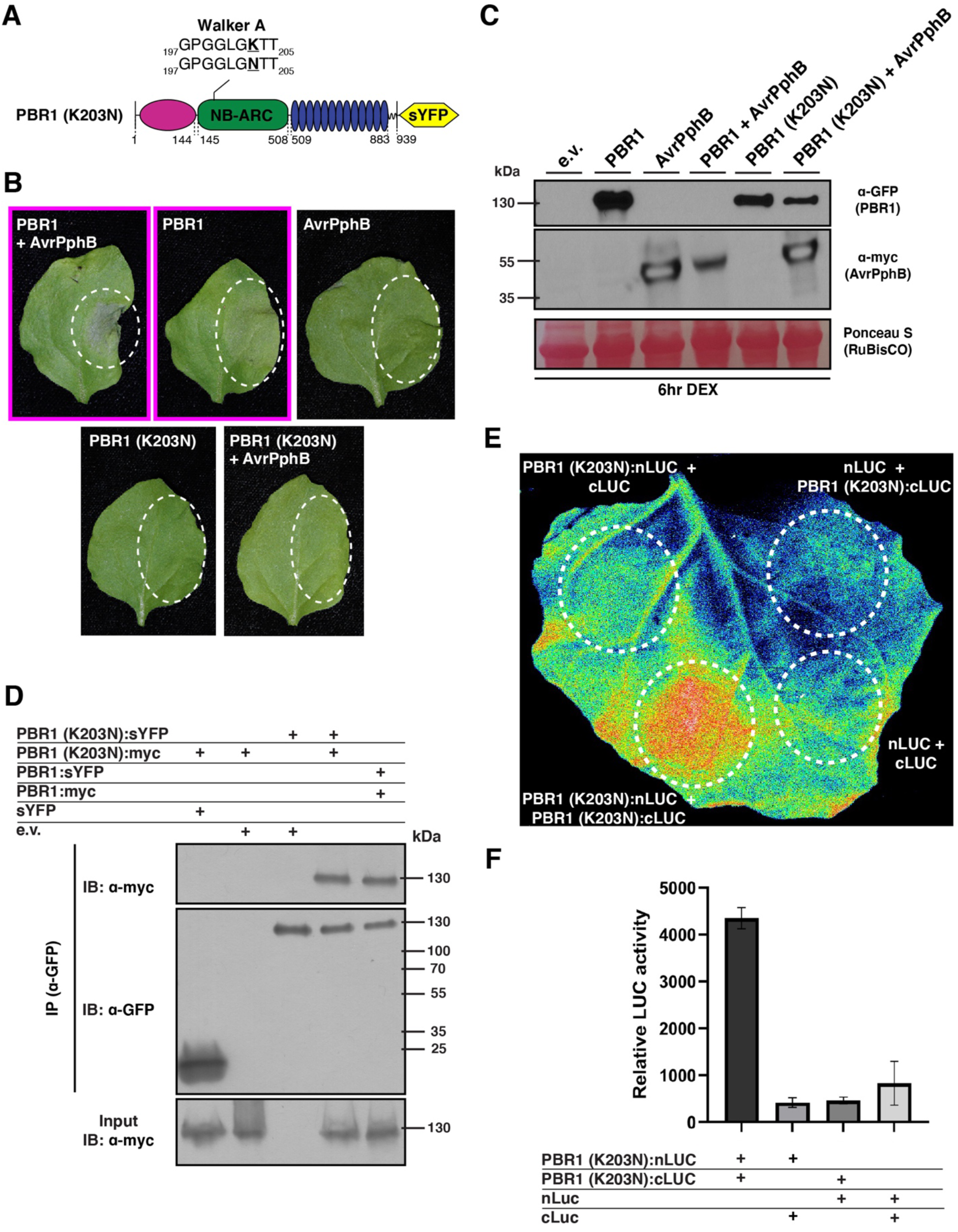
The K203N mutation within the NB-ARC domain of full-length PBR1 abolishes PBR1-mediated cell death but not self-association. **(A)** Schematic representation of the sYFP-tagged full-length PBR1 (K203N) derivative containing the K203N mutation. The P-loop/Walker A motif is shown and the K203N mutation is underlined. **(B)** The sYFP-tagged PBR1 (K203N) derivative fails to induce HR-like cell death when transiently expressed in *N. benthamiana*. Agroinfiltration was used to transiently co-express the indicated constructs into 3-week-old *N. benthamiana*. White circles indicate the agroinfiltration areas. Full-length PBR1:sYFP co-expressed with myc-tagged AvrPphB was used as a positive control. Leaves were harvested 24 hours post-transgene induction and a representative leaf was photographed under white light shortly thereafter. Photographs outlined with a magenta box indicate the agroinfiltrated leaf area showed observable cell death. **(C)** Immunoblot analyses showing protein accumulation of the PBR1 (K203N) derivative. The indicated constructs were transiently expressed in *N. benthamiana* using agroinfiltration. Total protein was extracted 6 hours following dexamethasone application. Equal amounts of total protein were resolved on 4-20% SDS-PAGE gels, blotted onto nitrocellulose membranes, and probed using either anti-GFP-horseradish peroxidase or anti-Myc-horseradish peroxidase antibodies. Ponceau staining of the RuBisCO large subunit was used as a loading control. **(D)** Co-immunoprecipitation (co-IP) assays were performed to detect the self-association of the PBR1 (K203N) derivative. The indicated constructs were transiently expressed in N. benthamiana using agroinfiltration and total protein was extracted 6 hours post-transgene induction. Protein samples were immunoprecipitated by GFP-Trap agarose bead slurry, followed by immunoblotting with the indicated antibodies. **(E and F)** SplitLUC assays show self-association of the PBR1 (K203N) derivative in *N. benthamiana*. PBR1 (K203N):nLUC and PBR1 (K203N):cLUC were coexpressed in *N. benthamiana* using agroinfiltration. Thirty hours following agroinfiltration, *N. benthamiana* leaves were sprayed with 1mM luciferin and a low-light cooled CCD imaging system was used to capture luciferase activity. Free cLUC coexpressed with PBR1 (K203N):nLUC, free nLUC coexpressed with PBR1 (K203N):cLUC, and free nLUC coexpressed with free cLUC were used as negative controls. Relative luciferase activity shown in panel E was quantified using a microplate luminescence reader (Infinite M200 Pro Plate Reader). Each data point consisted of three replicates and three independent experiments were performed.

Having shown PBR1 (K203N) failed to induce cell death, we next tested whether an intact P-loop/Walker A motif is required for the self-association of full-length PBR1. Notably, myc-tagged PBR1 (K203N) consistently co-immunoprecipitated with sYFP-tagged PBR1 (K203N) (Figure 7D). Similarly, split-luciferase complementation assays revealed detectable luciferase signals upon co-expression of PBR1 (K203N) fused to different halves of luciferase (Figure 7E and 6F). Collectively, these results indicate a functional P-loop/Walker A motif may not be required for the self-association of PBR1.

## DISCUSSION

We previously showed the NLR protein, PBR1, mediates recognition of AvrPphB protease activity, but it was unclear of the molecular mechanisms that lead to PBR1-dependent cell death. We, therefore, characterized PBR1 immune signaling in the absence and presence of AvrPphB using *Agrobacterium*-mediated transient expression in *Nicotiana benthamiana*. Here, we show that induction of cell death cannot be recapitulated by expressing the individual domains or truncations of PBR1, supporting the observation that PBR1 requires the cooperation of all domains *in cis*. Furthermore, we show that full length PBR1 is required for activation of cell death even when the autoactive mutant allele, PBR1 (D496V), is employed. Lastly, we show that PBR1 self-associates in the absence of AvrPphB, and such association is independent of a functional P-loop/Walker A motif.

PBR1-mediated induction of cell death is suppressed when an epitope tag is added to the N-terminus of both PBR1 and the PBR1 (D496V) derivative, but not the C-terminus (Figure 2). A similar loss-of-function in NLR-mediated immune signaling was reported for the immune receptors MLA1, MLA6, ZAR1, Sr35, and Pit when an epitope tag was fused to the N-terminus (Bieri et al., 2004; Bolus et al., 2020; Kawano et al., 2014; Wang et al., 2019a). Although our data are consistent with previous findings, it does not rule out the possibility that addition of an N-terminal epitope tag interferes with the subcellular localization of PBR1. For example, Kawano et al., (2014) showed that addition of an epitope tag to the N-terminus of the rice resistance protein, Pit, altered the localization from the plasma membrane to the cytosol. PBR1 is predicted to localize to the plasma membrane given that it recognizes plasma membrane-localized AvrPphB (Carter et al., 2019). Hence, future work should thus investigate the subcellular localization of both N-terminal and C-terminal-tagged PBR1.

The observation that none of the individual domains or fragments of PBR1 activate cell death when expressed individually or *in trans* is somewhat unique among CNLs. For example, expression of the CC–NB-ARC fragment from RPS2, RPS5, Sr33, Sr50, MLA10, Pvr4, and Rx consistently activated a robust cell death (Bai et al., 2012; Cesari et al., 2016; Day et al., 2005; Kim et al., 2018; Qi et al., 2012; Rairdan et al., 2008). Similarly, overexpression of the CC domain alone often induces an HR-like cell death response in *N. benthamiana* (Maekawa et al., 2011; Wang et al., 2015). Here, we show that regardless of whether the domains were expressed individually or *in trans* with or without AvrPphB, none of the constructs activated HR-like cell death, revealing that recapitulation of PBR1-mediated immune signaling requires the cooperation of all domains. However, it is a formal possibility that the absence of observed cell death is a result of differing subcellular localizations between full-length PBR1 and the individual domains *in cis*. Hence, future work should investigate the subcellular localization patterns of full-length PBR1 relative to that of the individual domains and truncations as such knowledge will be informative in determining the subcellular requirements for PBR1-mediated immune signaling.

Our data showing mutation of the aspartic acid within the MHD motif of full-length PBR1 activated elicitor-independent HR (Figure 4) is consistent with other CNLs (Bai et al., 2012; Bendahmane et al., 2002; Bolus et al., 2020; Chen et al., 2018; Gao et al., 2011; Kawano et al., 2014; Tameling et al., 2006). However, introducing the equivalent mutation within the NB-ARC domain, CC–NB-ARC, or NB-ARC–LRR truncations failed to activate HR-like cell death as strong as the PBR1 (D496V) derivative (Figure 4). Consistent with our observations, El Kasmi et al., (2017) showed that introducing the MHD to MHV mutation within full-length RPM1 activated HR-like cell death whereas the same mutation introduced into the NB-ARC domain, CC–NB-ARC, or NB-ARC–LRR truncations did not render these constructs autoactive. These data further support the notion that PBR1-mediated immune signaling requires the cooperation of all domains. Though we observed that wild-type PBR1 activated a mild cell death response by itself, PBR1 (D496V) consistently induced a stronger cell death response, supporting the notion that wild-type PBR1 protein is mainly in an inactive state. We predict that the mild cell death induced by wild-type PBR1 may be the result of the strong overexpression of PBR1 by the dexamethasone promoter, which may lead to improper inter-and intra-molecular associations.

Previous studies on N, RPS5, and Prf immune receptors indicate that their N-terminal domains provide a platform for self-association (Ade et al., 2007; Gutierrez et al., 2010; Mestre and Baulcombe, 2006). Consistent with other CNL proteins, we show that full-length PBR1 as well as the isolated domains and truncations self-associate in the absence of AvrPphB (Figure 5 and 6). Intriguingly, we showed that mutating the lysine residue within the P-loop/Walker A motif reduced PBR1-dependent cell death but did not abolish the self-association of full-length PBR1 (K203N) (Figure 7). Our data, therefore, suggests that activation, but not oligomerization, of PBR1 is dependent upon a functional P-loop/Walker A motif.

## ACKNOWLEDGEMENTS

The authors thank Ariel Sturgill-Helm for her excellent technical assistance and the Department of Horticulture and Landscape Architecture (Purdue University) for access to the low-light cooled CCD imaging system used in the split-luciferase assays. The authors would like to thank Andrew Read and Roger Innes for insightful discussions and critical reading of the manuscript. The funding bodies had no role in designing the experiments, collecting the data, or writing the manuscript. All opinions expressed in this paper are the author’s and do not necessarily reflect the policies and views of USDA, DOE, or ORAU/ORISE. USDA is an equal opportunity provider and employer.

## SUPPLEMENTARY FIGURE LEGENDS

**Supplemental Figure 1.**
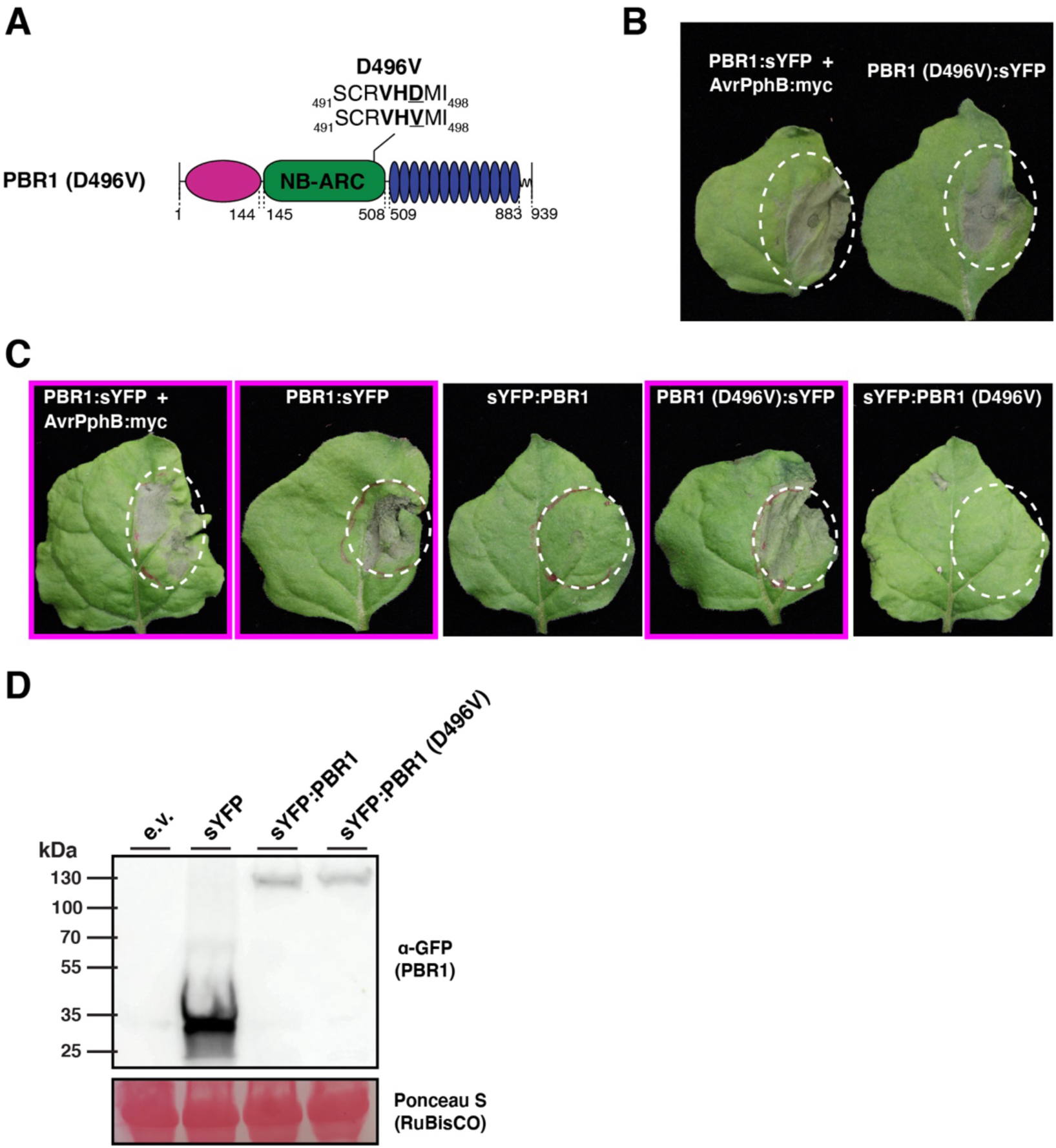
Fusing an epitope tag to the N-terminus of the auto-active PBR1 (D496V) derivative fails to activate cell death. **A)** Schematic illustration of the PBR1 (D496V) derivative. The MHD motif is shown in bold and the D496V mutation is underlined. **B)** Transient expression of the PBR1 (D496V) allele induces cell death. White circles indicate the agroinfiltration areas within the *N. benthamiana* leaves. Phenotypes were observed 24 hours following dexamethasone applications and a representative leaf was photographed under white light shortly thereafter. **C)** The N-terminal-tagged PBR1 (D496V) allele fails to induce HR-like cell death. Agroinfiltration was used to transiently express the indicated constructs into 3-week-old *N. benthamiana*. White circles indicate the agroinfiltration areas. Full-length PBR1:sYFP co-expressed with myc-tagged AvrPphB and PBR1 (D496V):sYFP were used as positive controls. Leaves were harvested 24 hours post-transgene induction and a representative leaf was photographed under white light shortly thereafter. Photographs outlined with a magenta box indicate the agroinfiltrated leaf area showed observable cell death. **D)** The sYFP:PBR1 (D496V) derivative expresses detectable protein. Immunoblot blot of total protein extracts from *N. benthamiana* leaf tissue agroinfiltrated with the indicated construct combinations. Total protein was extracted following 6 hours post-transgene induction and immunoblotted with anti-GFP antibodies. Ponceaus S staining solution was used as a control to show equal loading of total protein samples.

**Supplemental Figure 2.**
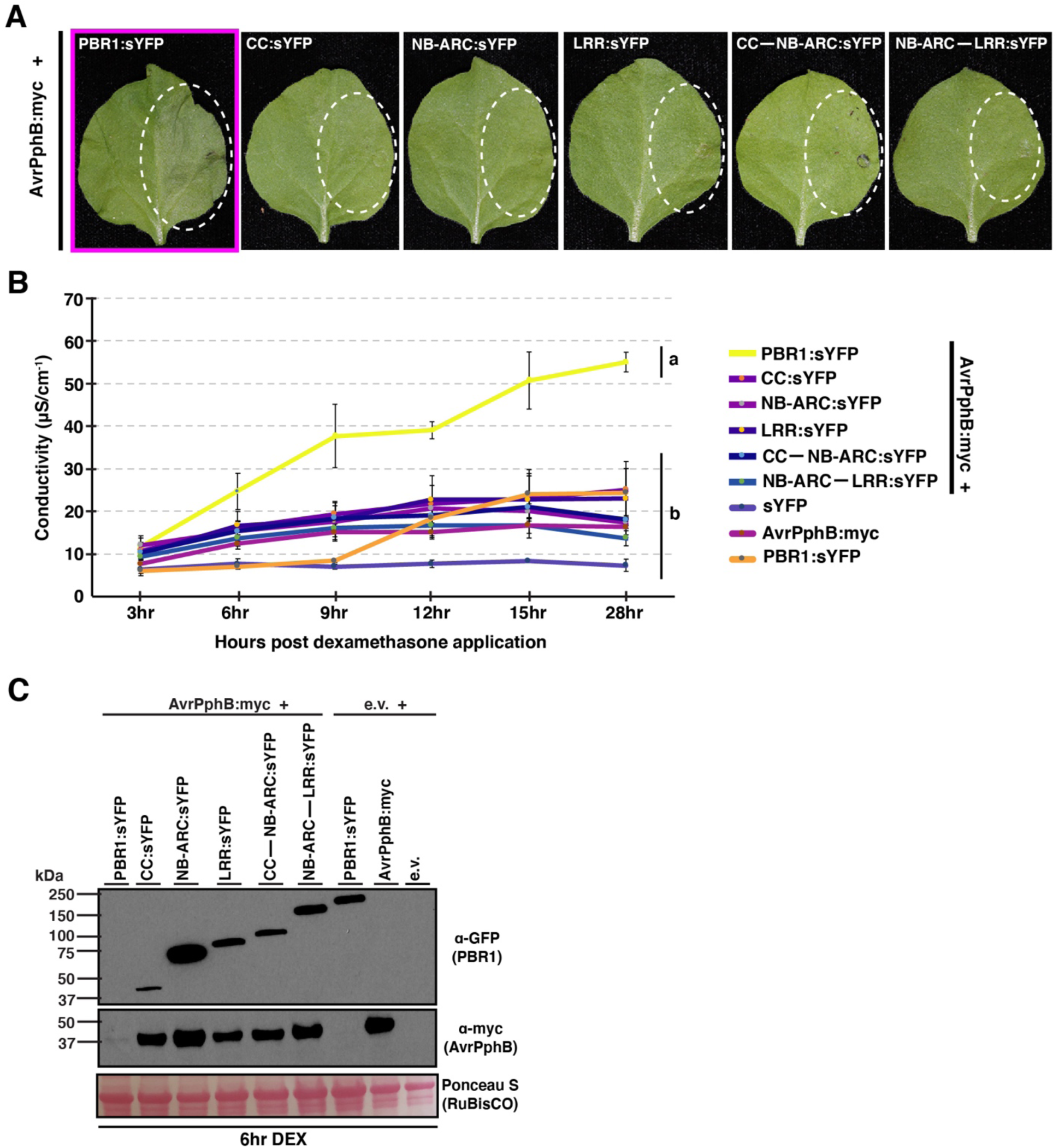
Lack of cell death induction by sYFP-tagged PBR1 domains and truncations when transiently co-expressed with AvrPphB in *N. benthamiana.* **(A)** Transient co-expression of truncation variants of PBR1 with AvrPphB do not induce an HR-like cell death response in *N. benthamiana*. The indicated constructs were transiently co-expressed in 3-week-old *N. benthamiana* using agroinfiltration. All proteins were under the control of a dexamethasone inducible promoter. PBR1:sYFP co-expressed with AvrPphB:myc was used as a positive control. White circles indicate the agroinfiltration areas within the *N. benthamiana* leaves. Phenotypes were observed 24 hours after dexamethasone induction and a representative leaf was photographed under white light shortly thereafter. Photographs outlined with a magenta box indicate the agroinfiltrated leaf area showed observable cell death. **(B)** Electrolyte leakage assay to quantify the cell death response. The indicated construct combinations were transiently co-expressed in *N. benthamiana* using agroinfiltration. All constructs were under the control of a dexamethasone inducible promoter. Conductivity was recorded at the indicated timepoints following transgene induction and is shown as mean SE (n = 4). Statistical significance was assessed using one-way ANOVA followed by Tukey-Kramer honestly significant difference analyses. Different letters represent significantly different means based on Tukey-Kramer honestly significant difference test (using α = 0.05). **(D)** Western blot of total protein extracts from *N. benthamiana* agroinfiltrated with the indicated construct combinations. Total protein was extracted following 6 hours post-transgene induction and immunoblotted with the indicated antibodies. Ponceaus S staining solution was used as a control to show equal loading of total protein samples. Expected M_r_s of proteins: PBR1:sYFP ∼133 kDa, CC:sYFP ∼43kDa, NB-ARC:sYFP ∼67kDa, LRR:sYFP ∼76kDa, CC—NB-ARC:sYFP ∼85kDa, and NB-ARC—LRR:sYFP ∼116kDa, AvrPphB:myc ∼33kDa.

**Supplemental Figure 3.**
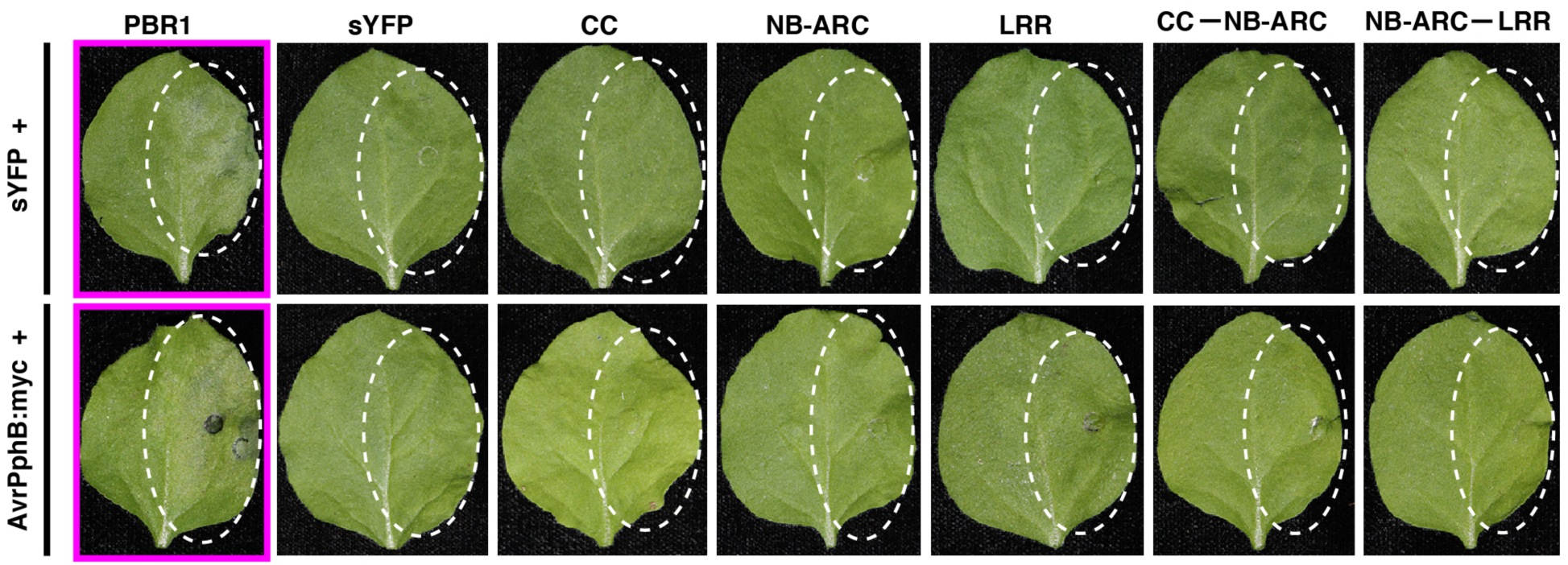
The presence of an epitope tag does not influence the lack of cell death induction by individual PBR1 domains and truncations co-expressed with or without myc-tagged AvrPphB. Non-epitope tagged PBR1 domains and truncations were co-expressed with either myc-tagged AvrPphB or sYFP in *N. benthamiana*. sYFP-tagged PBR1 co-expressed with AvrPphB-myc was used as a positive control. White circles indicate the agroinfiltration areas within the *N. benthamiana* leaves. Leaves were harvested 24 hours post-transgene induction and photographed under white light shortly thereafter. Photographs outlined with a magenta box indicate the agroinfiltrated leaf area showed observable cell death.

**Supplemental Figure 4.**
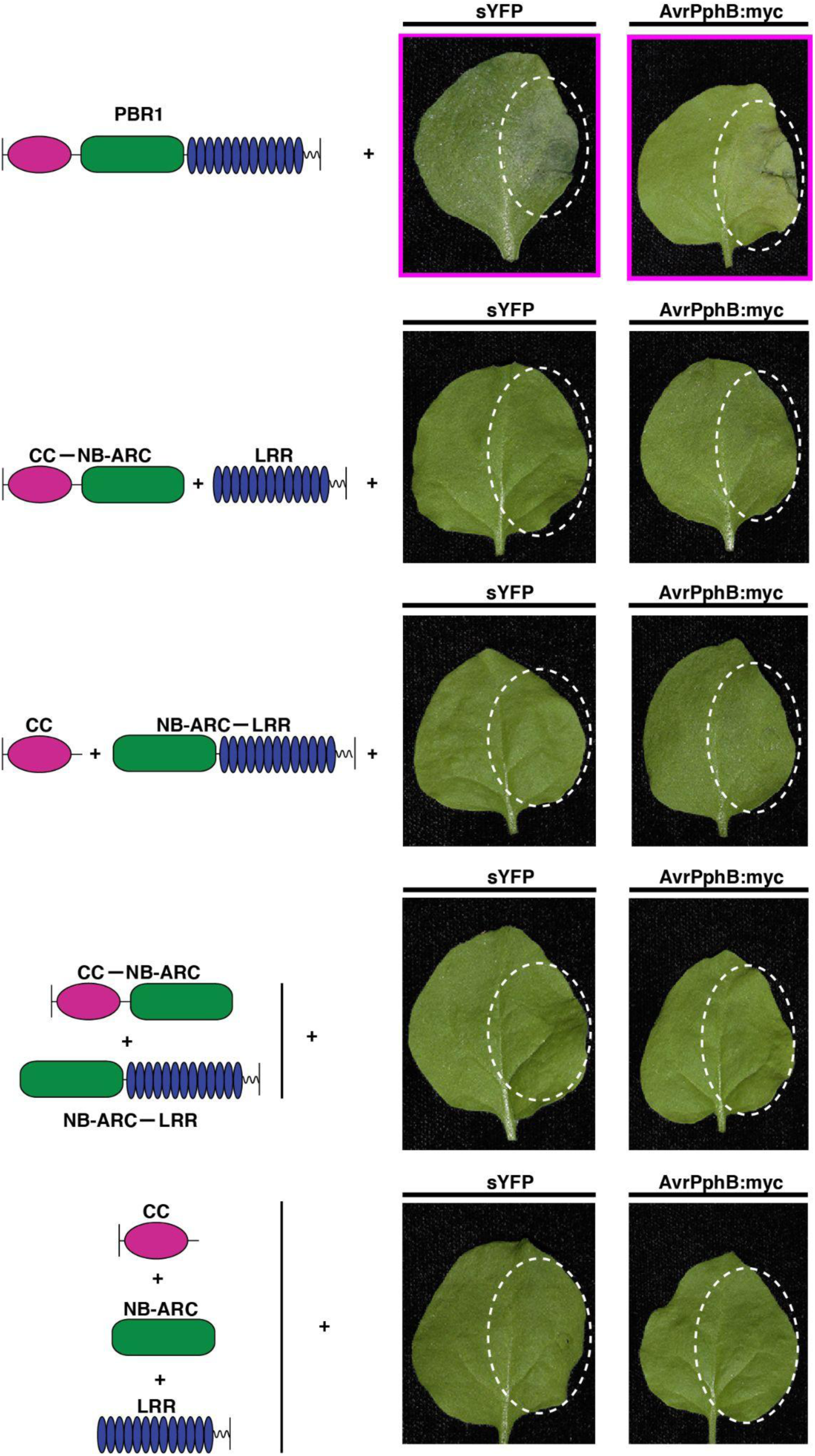
Transient co-expression of complementary PBR1 domains *in trans* with or without myc-tagged AvrPphB fail to activate cell death in *N. benthamiana*. Lack of cell death induction by complementary, non-epitope tagged PBR1 domains and fragments when expressed *in trans*. The indicated construct combinations were transiently co-expressed with either myc-tagged AvrPphB or sYFP into 3-week-old *N. benthamiana* using agroinfiltration. All proteins were under the control of a dexamethasone inducible promoter. Non-epitope tagged full-length PBR1 co-expressed with myc-tagged AvrPphB was used as a positive control. White circles indicate the agroinfiltration areas within the *N. benthamiana* leaves. Leaves were harvested 24 hours post-transgene induction and a representative leaf was photographed under white light shortly thereafter. Photographs outlined with a magenta box indicate the agroinfiltrated leaf area showed observable cell death.

**Supplemental Figure 5.**
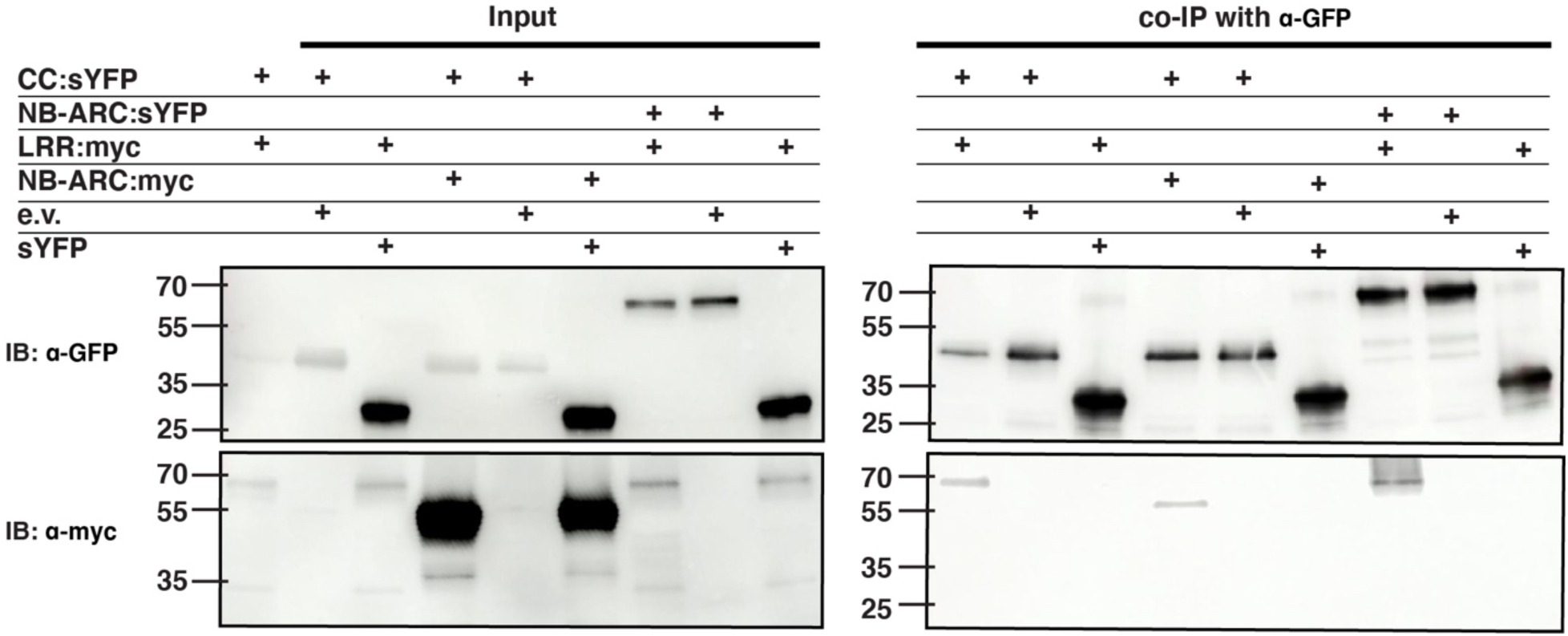
PBR1 domains show heterotypic associations in the absence of AvrPphB. The indicated constructs were transiently expressed in *N. benthamiana* using agroinfiltration and total protein was extracted 6 hours post-transgene induction. Protein samples were immunoprecipitated by GFP-Trap agarose bead slurry, followed by immunoblotting with the indicated antibodies.

